# Genetic Modification to Design a Stable Yeast-expressed Recombinant SARS-CoV-2 Receptor Binding Domain as a COVID-19 Vaccine Candidate

**DOI:** 10.1101/2020.11.09.373449

**Authors:** Wen-Hsiang Chen, Junfei Wei, Rakhi Tyagi Kundu, Rakesh Adhikari, Zhuyun Liu, Jungsoon Lee, Leroy Versteeg, Cristina Poveda, Brian Keegan, Maria Jose Villar, Ana C. de Araujo Leao, Joanne Altieri Rivera, Portia M. Gillespie, Jeroen Pollet, Ulrich Strych, Bin Zhan, Peter J. Hotez, Maria Elena Bottazzi

**Affiliations:** Texas Children’s Hospital Center for Vaccine Development, Houston, TX, USA; Departments of Pediatrics and Molecular Virology & Microbiology; National School of Tropical Medicine; Baylor College of Medicine, Houston, TX, USA; Department of Biology, Baylor University, Waco, TX, USA; James A. Baker III Institute for Public Policy, Rice University, Houston, TX, USA

**Author notes:** correspondence: Maria Elena Bottazzi, 1102 Bates St., Ste. 550 | Houston, TX 77030, Peter J Hotez, 1 Baylor Plaza, Houston TX 77030.

**Keywords:** coronavirus, *P. pastoris*, biophysical characterization, biotechnology

## Abstract

**Background:** Coronavirus disease 2019 (COVID-19) caused by SARS-CoV-2 has now spread worldwide to infect over 110 million people, with approximately 2.5 million reported deaths. A safe and effective vaccine remains urgently needed.

**Method:** We constructed three variants of the recombinant receptor-binding domain (RBD) of the SARS-CoV-2 spike (S) protein (residues 331-549) in yeast as follows: (1) a “wild type” RBD (RBD219-WT), (2) a deglycosylated form (RBD219-N1) by deleting the first N-glycosylation site, and (3) a combined deglycosylated and cysteine-mutagenized form (C538A-mutated variant (RBD219-N1C1)). We compared the expression yields, biophysical characteristics, and functionality of the proteins produced from these constructs.

**Results and conclusions:** These three recombinant RBDs showed similar secondary and tertiary structure thermal stability and had the same affinity to their receptor, angiotensin-converting enzyme 2 (ACE-2), suggesting that the selected deletion or mutations did not cause any significant structural changes or alteration of function. However, RBD219-N1C1 had a higher fermentation yield, was easier to purify, was not hyperglycosylated, and had a lower tendency to form oligomers, and thus was selected for further vaccine development and evaluation.

**General significance:** By genetic modification, we were able to design a better-controlled and more stable vaccine candidate, which is an essential and important criterion for any process and manufacturing of biologics or drugs for human use.

## 1. INTRODUCTION

Betacoronaviruses have caused major disease outbreaks in humans every decade since the beginning of the millennium. In 2002, severe acute respiratory syndrome coronavirus (SARS-CoV) was responsible for 8,098 infections and approximately 800 deaths. Beginning in 2012, the Middle East respiratory syndrome (MERS) caused by MERS-CoV has led to 2,519 infections and 866 associated deaths and the virus is still circulating in camels. First reported in China in December 2019, SARS-CoV-2, the pathogen that causes coronavirus disease 2019 (COVID-19), has now rapidly spread worldwide and infected more than 110 million people and caused approximately 2.5 million deaths up to early March 2021 [1].

A coronavirus virion consists of membrane, envelope, nucleocapsid, and spike (S) proteins. Of these, the S proteins are commonly selected as vaccine candidates due to their critical functions in host cell entry [2-7]. However, even though no preclinical or human trials have yet reported that immunopathology could be triggered by the full-length S protein of SARS-CoV-2 after viral challenge [8], previous preclinical studies in mice indicated that the full-length SARS-CoV S protein could induce eosinophilic immune enhancement [9, 10] and lung immunopathology [11, 12], possibly due to epitopes outside of the receptor-binding domain (RBD). The SARS-CoV RBD vaccine was shown to induce high levels of virus-neutralizing antibodies, minimized or abrogated eosinophilic immune enhancement in mice compared to the full-length S protein, and can be easily scaled for production [12, 13]; therefore, some groups consider the RBD an attractive vaccine immunogen alternative to using the full-length S protein [14-19]. Beyond our findings with SARS-CoV, additional studies have already shown that mammalian cell-expressed SARS-CoV-2 RBD triggered high specific total IgG as well as neutralizing antibody titers in mice [7] and that the vaccination of insect cell-expressed RBD protected non-human primates (NHPs) from SARS-CoV-2 challenge *in vivo* with no significant lung histopathological changes, further supporting a rationale for using the RBD as antigen to develop an efficacious vaccine [20].

Based on previous experience in developing the SARS-CoV RBD vaccine [4, 13, 17] and the high amino acid sequence similarity between SARS-CoV and SARS-CoV-2 spike proteins [21-23], we selected a 219 amino-acid sequence of SARS-CoV-2 (RBD219-WT), spanning residues 331-549 of the S protein [4]. The sequence includes two glycosylation sites at N331 and N343 [24]. However, hyperglycosylation as well as dimer formation were observed during the production of RBD219-WT. The variable lengths of the glycans attached to RBD219-WT posed a challenge to the purification process and no uniform band was observed by SDS-PAGE. In addition, we observed dimerization of the antigen that might further impact the quality of antigen. To resolve these issues, we first removed Asn331 from the sequence, generating RBD219-N1, and then also mutated C538 to alanine (RBD219-N1C1). RBD219-N1C1 constitutes a better-controlled and more stable protein, which is an important and essential criterion for any process and manufacturing of a biologic for human use. The immunogenicity of these vaccine candidates was recently tested, demonstrating that when formulated with Alhydrogel®, these yeast-expressed RBDs triggered equivalent high titers of neutralizing antibodies [25].

Here, we examined the production yield of three tag-free RBDs in the fermentation supernatant prior to purification. Two of the RBD constructs resulted in high fermentation yields, however, for the RBD219-N1 construct, we only achieved low fermentation yields with likely high host cell protein levels. This impeded us from getting any purified RBD219-N1 protein, hence, a hexahistidine tagged construct (RBD219-N1+His) was created to obtain purified protein for further characterization. Biophysical characteristics, and *in vitro* functionality of these three purified recombinant RBD proteins were evaluated. Based on the results, we identified the RBD219-N1C1 construct best suited for advancement. This construct and its initial process for protein production have been transferred to an industrial manufacturer, who has successfully produced and scaled it and has now advanced this vaccine candidate into a Phase I/2 clinical trial [26].

## 2. MATERIALS AND METHODS

### 2.1. Cloning and expression of SARS-CoV-2 RBDs in yeast *Pichia pastoris*

The DNAs encoding RBD219-WT (residues 331–549 of the SARS-CoV-2 spike protein, GenBank: QHD43416.1), RBD219-N1 (residues 332-549), and RBD219-N1C1 (residues 332-549, C538A) were codon-optimized based on yeast codon usage preference and synthesized by GenScript (Piscataway, NJ, USA), followed by subcloning into the *Pichia* secretory expression vector pPICZαA (Invitrogen) using EcoRI/XbaI restriction sites. A hexahistidine tag version for RBD219-WT and RBD219-N1 (namely, RBD219-WT+His and RBD219-N1+His, respectively) was also generated by adding additional DNA encoding six histidine residues at the C-terminus to facilitate purification as a backup. The recombinant plasmid DNAs were then transformed into *Pichia pastoris* X-33 by electroporation. The expression of the recombinant RBDs was confirmed by induction with 0.5% methanol at 30 °C for 72 hours. The seed stock in 20% glycerol of each recombinant construct was then generated as described previously [13].

### 2.2. Fermentation and purification of SARS-CoV-2 RBDs

RBD219-WT, RBD219-N1, and RBD219-N1C1 in pPICZαA/*P. pastoris X33* clones were fermented in 5 L vessels as described previously with minor modifications [13]. Briefly, the seed stock of each construct was used to inoculate 0.5 L Buffered Minimal Glycerol (BMG) medium until the OD_600_ reached 10±5.

Depending on the OD_600_ of overnight culture, 86-270 mL of the culture was then used to inoculate 2.5 L sterile low salt medium (LS) in the fermenter containing 3.5 mL/L PTM1 trace elements and 3.5 mL/L 0.02% d-Biotin to reach to initial OD of 0.5. Fermentation was initiated at 30 °C and pH 5.0, while the gas and agitation were adjusted to maintain dissolved oxygen (DO) at 30%. Upon DO spike, the pH was ramped up to 6.5 using 14% ammonium hydroxide, and the temperature was lowered to 25°C over 1 hour and methanol was then pumped in from 0.8 mL/L/h to 11 mL/L/h and the pH was adjusted to 6.0 using 14% ammonium hydroxide over 6–8 hours. Induction was maintained at 25 °C with minor methanol feed adjustments, as needed, for 70 hours. After fermentation, the culture was harvested by centrifugation. The fermentation supernatant (FS) was then evaluated by SDS-PAGE and Western blot.

To purify RBD219-WT, the FS was first filtered through a 0.45 µm filter followed by a negative capture step with a Q Sepharose XL (QXL) column in 30 mM Tris-HCl, pH 8.0 to remove some host cell protein. The flow-through from the QXL column was then further purified by a Butyl Sepharose HP column and a Superdex 75 size exclusion column (SEC). Due to the low target protein yield and large amounts of impurities present in RBD219-N1 fermentation, we were unable to successfully purify the tag-free RBD219-N1 by the same approach as RBD219-WT. Instead, we purified the using the hexahistidine tagged version (RBD219-N1+His, where six additional histidine residues were expressed at the C-terminus of RBD219-N1); to purify RBD219-N1+His, HisTrap immobilized metal affinity column was used followed by Superdex 75 chromatography. Finally, to purify RBD219-N1C1, the FS was filtered through a 0.45 µm filter before a Butyl Sepharose HP column followed by a Superdex 75 column. The final buffer for these three proteins was TBS (20 mM Tris, 150 mM NaCl, pH 7.5).

### 2.3. SDS-PAGE and Western Blot

RBD219-WT, RBD219-N1+His, and RBD219-N1C1 were loaded on 4-20% Tris-glycine gels, and stained with Coomassie Blue or transferred to a polyvinylidene difluoride membrane and probed with a monoclonal anti-SARS-CoV-2 Spike rabbit antibody recognizing the RBD region (Sino Biological, Beijing, China; Cat # 40150-R007) to evaluate the size and confirm the identity. These three RBDs were also treated with PNGase-F (New England Biolabs, Ipswich, MA, USA; Cat# P0704S) following the manufacturer’s instruction and loaded onto SDS-PAGE gels to evaluate the impact of size caused by glycosylation. Western blotting was also used to evaluate the fermentation yield; in short, serially diluted purified RBD protein corresponding to the construct in the fermentation run was loaded on the Tris-glycine gels with a fixed volume of undiluted fermentation supernatant of different RBD constructs.

A log-log plot of RBD intensity versus the known amount of loaded RBD was graphed and the linear regression was calculated from the plot.

### 2.4. Size and Protein Aggregation Assessment by Dynamic Light Scattering

Purified RBDs were adjusted to 1 mg/mL in TBS in three to four replicates to evaluate the hydrodynamic radius and molecular weight using a DynaPro Plate Reader II (Wyatt Technology) based on a globular protein model. The sizes of these RBDs at room temperature were monitored for approximately 30 days. Additionally, to evaluate the tendency of protein oligomerization among different RBDs, these purified proteins were concentrated to approximately 7.5 mg/mL and serially diluted to approximately 0.66 mg/mL to calculate the diffusion interaction parameter (k_D_) for each RBD using the following equation [13, 27]:

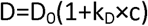

where D is the measured diffusion coefficient, D_0_ is the coefficient of the RBDs at an infinite dilution, and k_D_ is the diffusion interaction parameter.

### 2.5. Hydrophobicity Assessment by Extrinsic Fluorescence

Purified RBDs and two controls (BSA and lysozyme) at concentrations of 7.8 ng/mL - 1.0 mg/mL in TBS were probed with 12.5 nM Nile Red (Sigma-Aldrich, St. Louis, MO, USA; Cat # 3013) to generate extrinsic fluorescence for surface hydrophobicity measurement, as described previously [13, 28]. The emission spectra were recorded from 550 to 700 nm with the gain set at 130 after excitation at 520 nm using a BioTek Synergy H4 plate reader. The surface hydrophobicity of RBD219-WT, RBD219-N1+His, RBD219-N1C1, BSA, and lysozyme was determined using the slope of the emission peak intensity versus protein concentration plot, as described previously [13, 29].

### 2.6. Structural Assessment by Circular Dichroism (CD)

Purified RBDs were diluted with deionized water to a final concentration of 0.2 mg/mL and loaded in a 0.1 cm path cuvette. Please note that dilution with water was to reduce the chloride ion content which is known to interfere with the CD absorbance. CD spectra were obtained from 250 to 190 nm with a Jasco J-1500 spectrophotometer set at 100 nm/min and a response time of 1 s at 25°C. The obtained CD data were analyzed using a CD Analysis and Plotting Tool (https://capito.uni-jena.de/index.php). In addition, the RBDs (0.5 mg/mL) were heated from 25 °C to 95 °C for a denaturation profile analysis.

### 2.7. Structural Assessment by Thermal shift

To evaluate the thermal stability of the tertiary structures for these three RBDs, a 1 mg/mL solution of these proteins was mixed with the Protein Thermal Shift™ Dye kit (Thermo Fisher, Waltham, MA, USA; Cat # 4461146) in triplicate as per the manufacturer’s instructions. Briefly, 5 µL of Protein Thermal Shift buffer was mixed with 12.5 µL of 1.0 mg/mL RBD and 2.5 µL of 8x Protein Thermal Shift dye in each PCR tube in three to four replicates. These PCR tubes were vortexed briefly and centrifuged at 1000x g for 1 minute to remove any bubbles. The samples were further heated from 25 °C to 95 °C to monitor the fluorescence intensity change using a ViiA™ 7 Real-Time PCR system.

### 2.8. *in vitro* Functionality Assay by ELISA (ACE-2 binding)

Ninety-six-well ELISA plates were coated with 100 µL 2 µg/mL RBD219-WT, RBD219-N1+His, RBD219-N1C1, or BSA overnight in triplicate at 4°C followed by blocking with PBST/0.1% BSA. Once the plates were blocked, 100 µL serially diluted ACE-2-hFc (LakePharma, San Carlos, CA, USA; Cat # 46672) with an initial concentration of 50 µg/mL was added to the wells. The plates were incubated at room temperature for 2 hours to allow ACE-2 to interact with each RBD. After this binding step, the plates were washed with PBST four times followed by adding 100 µL 1:10,000 diluted HRP conjugated anti-human IgG antibodies (GenScript, Piscataway, NJ, USA; Cat # A00166) and incubating for 1 hour at room temperature. Finally, 100 µL TMB substrate was added and incubated for 4 minutes in the dark to react with HRP. The reaction was terminated with 100 µL HCl and absorption readings were taken at 450 nm using a BioTek EPOCH 2 microplate reader. The results were analyzed using one-way ANOVA to evaluate the statistical difference among the ELISA data generated for these three RBDs.

## 3. RESULTS

### 3.1. Cloning, expression yield, and protein integrity

The SARS-CoV-2 RBD vaccine antigen was originally designed and expressed using the same approach as reported previously for the 70% homologous SARS-CoV RBD equivalent [13]. Three recombinant constructs, RBD219-WT, RBD219-N1, and RBD219-N1C1 (Figure 1), were transformed into *P. pastoris* and expressed at the 5 L fermentation scale.

**Figure 1.**
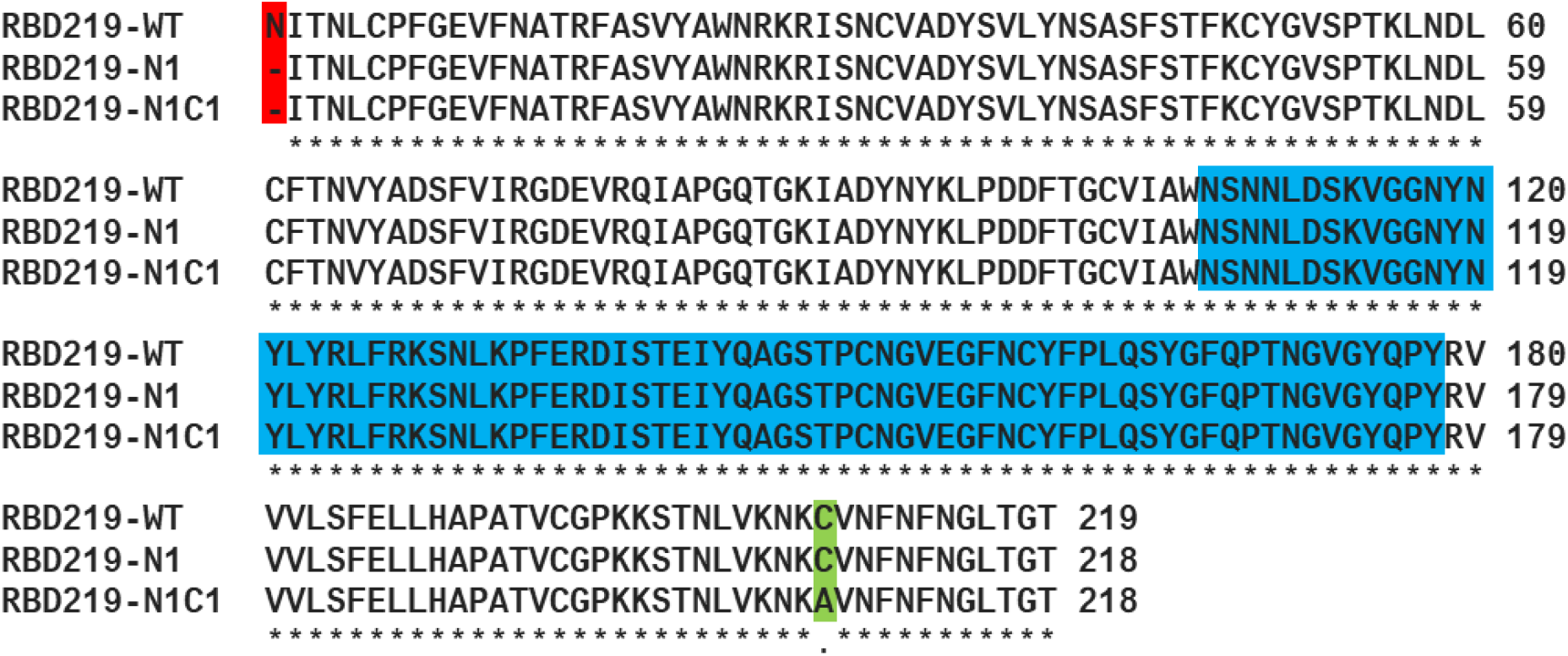
Sequence comparison among RBD219-WT, RBD219-N1, and RBD219-N1C1. The deleted asparagine in the first glycosylation site is highlighted in red while the mutated cysteine (to alanine) is highlighted in green. The receptor-binding motif is highlighted in blue [21, 56].

The harvested fermentation supernatants (FS) of RBD219-WT, RBD219-N1, and RBD219-N1C1 were first analyzed by SDS-PAGE (Figure 2A) and their fermentation yields were quantified using western blot (Supplementary Figure S1 and Table 1). When observing the FS profile for RBD219-WT on SDS-PAGE gel (Figure 2A) and western blot membrane (Supplementary Figures S1A and S1C), large amounts of protein bands were not recognized by the anti-SARS-CoV-2 Spike rabbit antibody, possibly indicating the presence of host cell proteins (HCP), as well as yeast-derived hyperglycosylation. The fermentation yield of RBD219-WT was estimated as 142 ± 8 mg/L of fermentation supernatant (Table 1).

**Figure 2.**
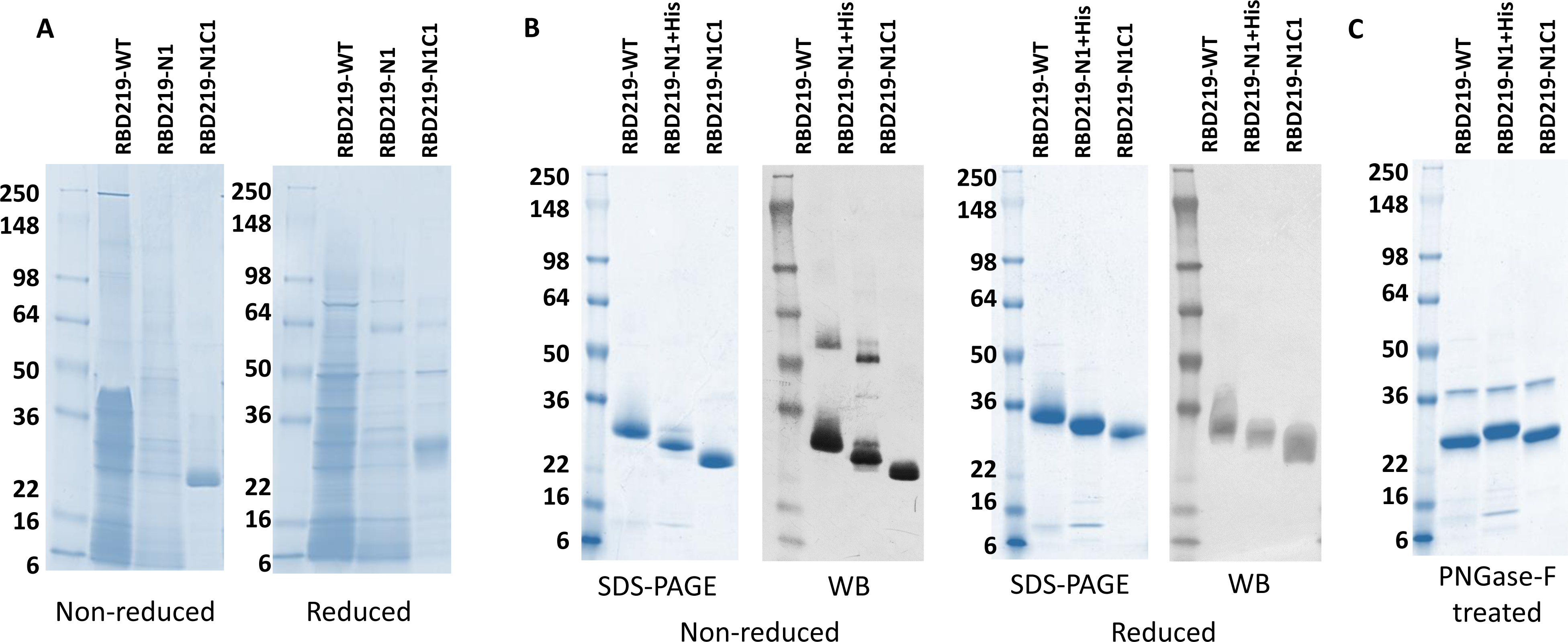
Coomassie Blue stained SDS-PAGE and western blot probed with anti-SARS-CoV-2 Spike rabbit antibody. (A) SDS-PAGE gel of 10 µL fermentation supernatant for tagged-free RBD219-WT, RBD219-N1, and RBD219-N1C1; (B) Coomassie Blue stained SDS-PAGE gel of 3 µg purified RBDs or western blot of 1.5 µg of the purified RBDs under non-reduced and reduced conditions. (C) SDS-PAGE of 3 µg PNGase-F treated purified RBDs; please note that the 37 kDa band observed on the PNGase-F treated gel is the N-glycosidase PNGase-F enzyme.

**Table 1.** Assessment of the fermentation supernatants for clones RBD219-219, RBD219-N1, and RBD219-N1C1. *Fermentation yield was measured by probing the RBD with the same specific antibody followed by densitometry (Supplementary Figure S1). FS: fermentation supernatant; WB: western blot.

| Molecule | Fermentation yield*<br>(mg RBD/L of FS) | Level of<br>Impurity<br>on SDS-PAGE | Hyperglycosylation on<br>WB | Dimer formation on<br>WB |
| --- | --- | --- | --- | --- |
| RBD219-WT | 142 ± 8 | High | Yes | Yes |
| RBD219-N1 | 50 ± 13 | Mid | No | Yes |
| RBD219-N1C1 | 280 ± 70 | Low | No | Yes |

To address the yeast-derived hyperglycosylation, we cloned and expressed RBD219-N1, in which the N-terminal asparagine residue was removed. However, the fermentation yield of RBD219-N1 was low (50 ± 13 mg/L of FS, Table 1), and the level of non-RBD proteins, likely HCP impurities, remained a concern (Figure 2A and Supplementary Figures S1E and S1F). Due to the low yield and high level of non-target-specific proteins, we were unable to purify tag-free RBD219-N1; thus, the hexahistidine tagged construct (RBD219-N1+His) was used to obtain purified protein for later characterization. We also observed that both RBD219-WT and RBD219-N1 formed dimers during fermentation. This suggested the potential formation of an intermolecular disulfide bond between a potential free cysteine found in the molecule, and therefore, a C538A-mutated form of RBD219-N1, RBD219-N1C1, was constructed. The rationale is that there are nine cysteine residues in the RBD, where we assumed that eight residues formed intramolecular bonds, leaving the last cysteine available for undesired intermolecular cross-linking. The new C538A-mutated construct based on the RBD219-N1 backbone was able to express the protein (RBD219-N1C1) with low yeast-derived hyperglycosylation, without the presence of extensive non-RBD specific proteins or HCPs (Figure 2A), and with a fermentation yield of 280 ± 70 mg/L of FS (Table 1). Even though, when fermentation yields were analyzed using western blot, one could observe a dimer form of RBD219-N1C1 in the fermentation supernatant (Supplementary Figures S1I and S1K), such dimer was not seen after purification in the final purified protein (Figure 2B). After treating the three purified RBDs with PNGase-F to remove N-glycans, RBDs with similar size were observed (Figure 2C; Please note that the band at 37 kDa was PNGase-F enzyme), which confirmed that the size differences were likely due to the yeast-derived glycosylation.

### 3.2. Size evaluation and protein-interaction assessment by DLS

Dynamic light scattering (DLS) was used to evaluate the size of the purified RBDs in solution. The results indicated that SARS-CoV-2 RBD219-WT has a slightly larger hydrodynamic radius (2.79 ± 0.01 nm) than RBD219-N1C1 (2.56 ± 0.00 nm), and thus a higher calculated molecular weight (37.5 ± 0.5 kDa) than RBD219-N1C1 (30 ± 0.0 kDa) (Figure 3A), presumably due to the additional N-glycans. Interestingly, even though RBD219-N1+His was less glycosylated due to the removal of the first glycosylation site, its hydrodynamic radius (2.74 ± 0.01 nm, 35.7 ± 0.5 kDa) was similar to RBD219-WT; thus, it was suspected that potential oligomer formation might have occurred. We further determined the diffusion interaction parameter (K_D_) of three RBDs (Figure 3B and Table 2) to evaluate the level of protein-protein interaction. The results indicated that all three RBDs in TBS showed negative K_D_ values, implying protein attractions. However, it was noticed that the K_D_ of RBD219-N1C1 (−16.3 mL/g) was similar to that of RBD219-WT (−14.9 mL/g), while that of RBD219-N1+His was almost double (−29.7 mL/g), suggesting that RBD219-N1+His was even more prone to form oligomers. Additionally, when we monitored the changes of the molecular weight for RBDs stored at room temperature, it was observed that both the molecular weight of RBD219-WT and RBD219-N1+His increased over time while the size of RBD219-N1C1 remained unchanged at ∼30kDa (Figure 3C).

**Figure 3.**
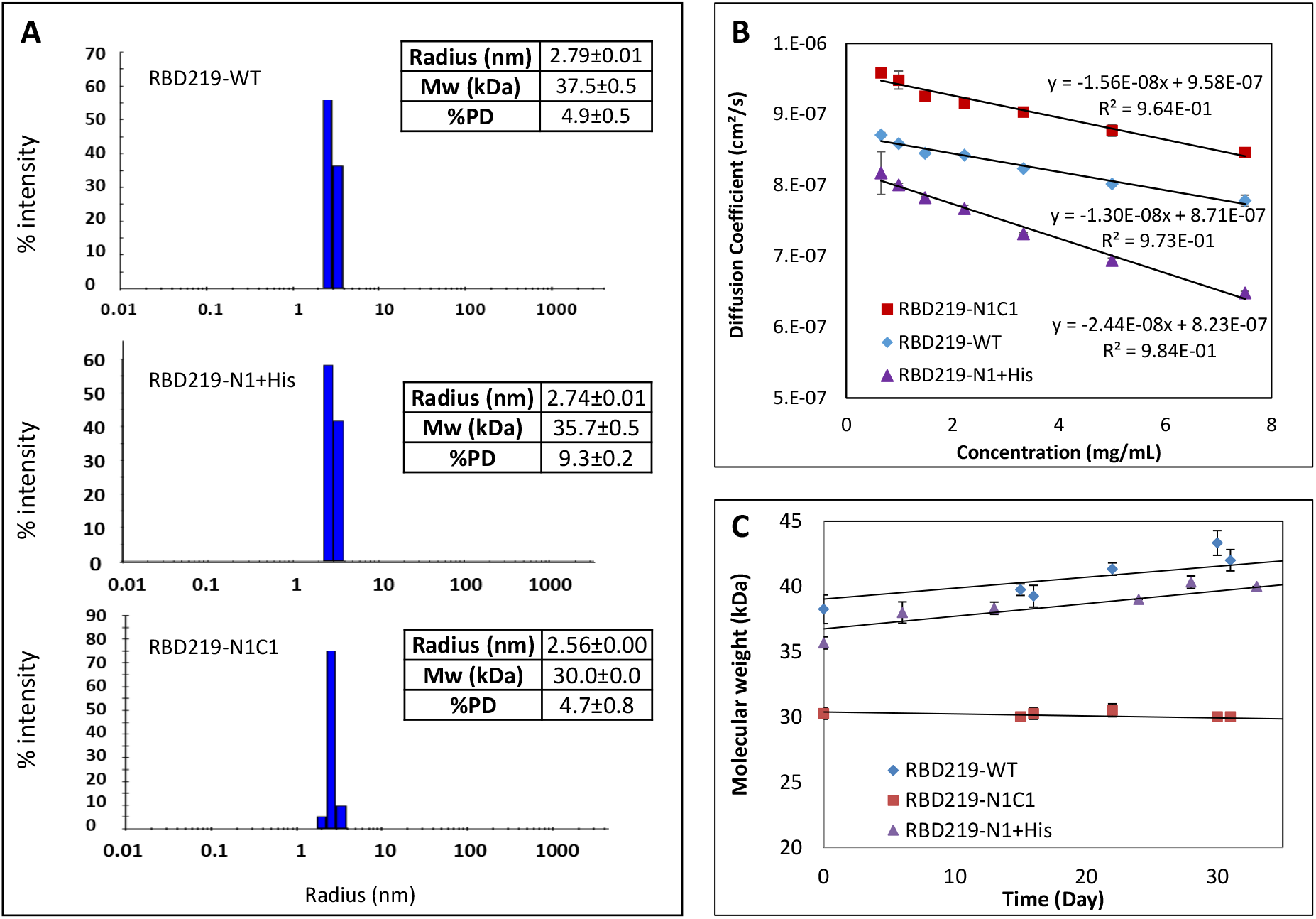
Dynamic light scattering results for RBD219-WT, RBD219-N1+His, and RBD219-N1C1. (A) Measured Stokes radii and molecular weights. (B) Diffusion coefficient vs. concentration plot to evaluate the diffusion interaction parameter (C) Stability study to monitor the changes of the molecular weight.

**Table 2.** Aggregation assessment by diffusion interaction parameter (K_D_) using DLS and hydrophobicity assessment by extrinsic fluorescence.

| Protein | Dynamic light scattering | Extrinsic fluorescence |  |
| --- | --- | --- | --- |
| | Diffusion interaction<br>parameter $K_D$ (mL/g) | Peak wavelength<br>$\lambda_{max}$ (nm) | Surface hydrophobicity<br>$S_0$ (RFU*L/mg) |
| RBD219-WT | -14.9 | 658 | 3.1 |
| RBD219-N1+His | -29.7 | 642 | 9.8 |
| RBD219-N1C1 | -16.3 | 658 | 2.8 |
| BSA | N/A | 634 | 53.4 |
| Lysozyme | N/A | 662 | 1.1 |

### 3.3. Hydrophobicity Assessment Using Extrinsic Fluorescence

In this study, we used Nile red dye to probe these three purified RBDs to further evaluate their surface hydrophobicity in TBS (Figures 4A and 4B). In general, a blue shift in emission peak wavelength λmax, or a lower λmax value, indicates an increase in hydrophobicity due to more binding of Nile red to the protein. The emission peaks λmax of RBD219-WT, RBD219-N1+His, and RBD219-N1C1 in TBS were determined as 658, 642 and 658 nm, respectively, indicating that RBD219-N1+His was more hydrophobic than the other two RBD molecules, and these three RBDs were less hydrophobic than BSA (λmax = 634 nm) and more hydrophobic than lysozyme (λmax = 662 nm) (Figure 4A and Table 2). Additionally, the surface hydrophobicity (S_0_) calculated by the plot of fluorescence intensity vs concentration (Figure 4B and Table 2) indicated that RBD219-WT and RBD219-N1C1 shared similar surface hydrophobicity (S_0; RBD219-WT_ = 3.14 RFU*L/mg and S_0; RBD219-N1C1_= 2.78 RFU*L/mg, respectively), while RBD219-N1+His showed higher surface hydrophobicity (S_0; RBD219-N1+His_ = 9.82 RFU*L/mg). Overall, the results indicated S_0; BSA_ > S_0; RBD219-N1+His_ > S_0; RBD219-WT_ ∼ S_0; RBD219-N1C1_> S_0; Lysozyme_) which were consistent with the observation based on the wavelength of the extrinsic fluorescence emission peak.

**Figure 4.**
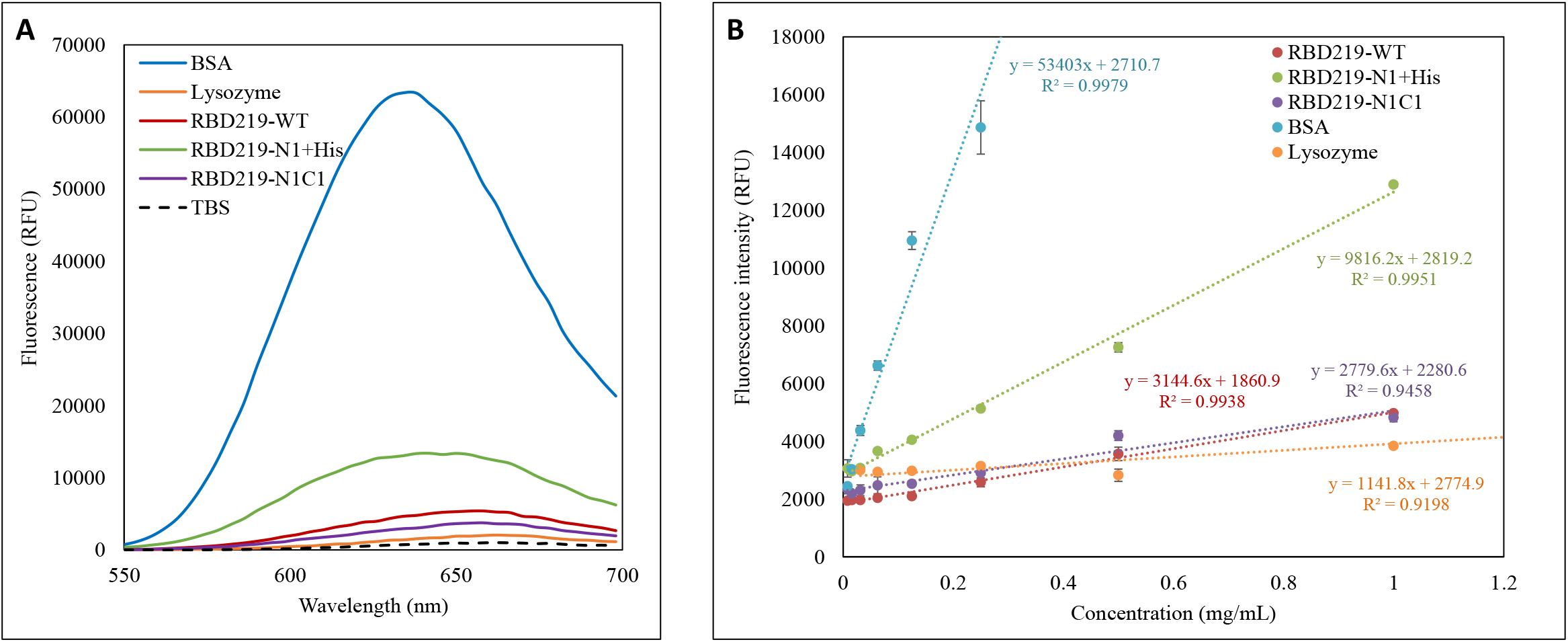
Extrinsic fluorescence results for RBD219-WT, RBD219-N1+His, and RBD219-N1C1. (A) Excitation wavelength scan to obtain the peak emission wavelength λmax; (B) Fluorescence intensity vs concentration plot to evaluate surface hydrophobicity.

### 3.4. Secondary structure thermal stability assessment

When far-UV CD spectrometry was performed to investigate the secondary structure of the three purified RBDs, we observed a very similar secondary structure with 8% alpha-helix and 33% beta-sheet (Fig. 5A). The thermal stability of the secondary structures was evaluated by heating the samples from 25 °C to 95 °C (Figs. 5B-5D) and CD melting curves and their derivatives were further examined at 231 nm (Figures. 5E-G). Based on the derivative, the average melting temperatures (Tm) were 52.3 °C, 53.3 °C, and 51.9 °C for RBD219-WT, RBD219-N1+His, and RBD219-N1C1 respectively with only 1.1% of the variation of coefficient (%CV) among these melting temperatures, suggesting similar thermal stability.

**Figure 5.**
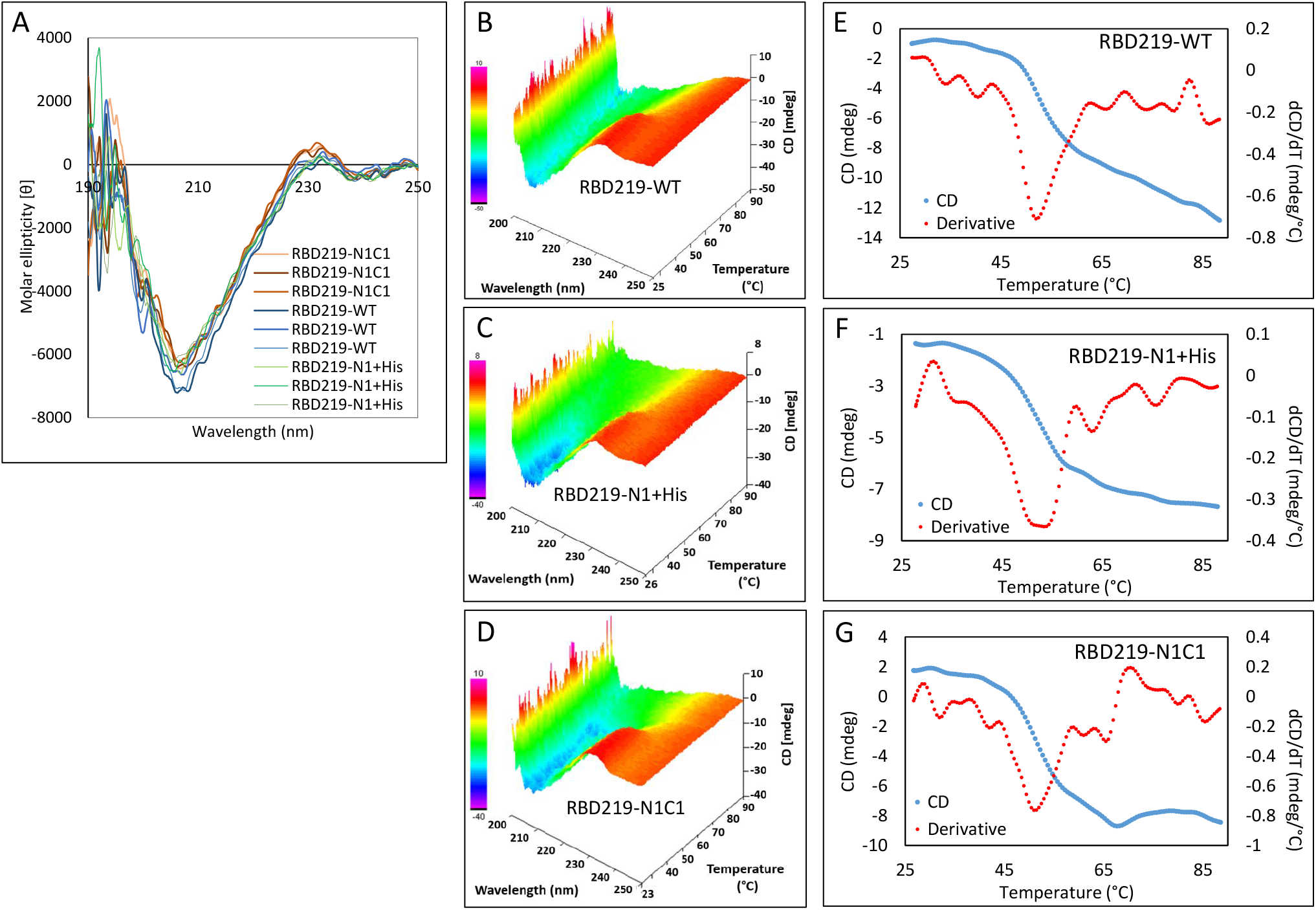
Circular dichroism data for RBD219-WT, RBD219-N1+His, and RBD219-N1C1. (A) Circular dichroism spectra; Thermal map of circular dichroism spectra for (B) RBD219-WT, (C) RBD219-N1+His, and (D) RBD219-N1C1. Melting profile of (E) RBD219-WT, (F) RBD219-N1+His, and (G) RBD219-N1C1.

### 3.5. Tertiary structure thermal stability assessment

Thermal shift assays were used for evaluating the thermal stability of the tertiary structure for the three purified RBD proteins. The melting curve (Figure 6A) indicated that the onset temperature (T_on_) for these RBDs was approximately 38 °C, and the higher initial fluorescence of RBD219-N1+His observed in Figure 6A implied that more thermal shift dye bound to the molecule, which suggested higher hydrophobicity of this molecule than for the other two RBDs. This finding is consistent with the observation in the extrinsic fluorescence study. The derivatives indicated the melting temperatures (T_m_) for RBD219-WT, RBD219-N1+His, and RBD219-N1C1 as 50.6 ± 0.5 °C, 49.2 ± 0.5 °C and 50.8 ± 0.4 °C, respectively (Figure 6B) with a %CV value of 1.4%. The similar thermal shift profile and thermal stability observed in the thermal shift assays further suggested that the three RBDs likely shared similar tertiary structures.

**Figure 6.**
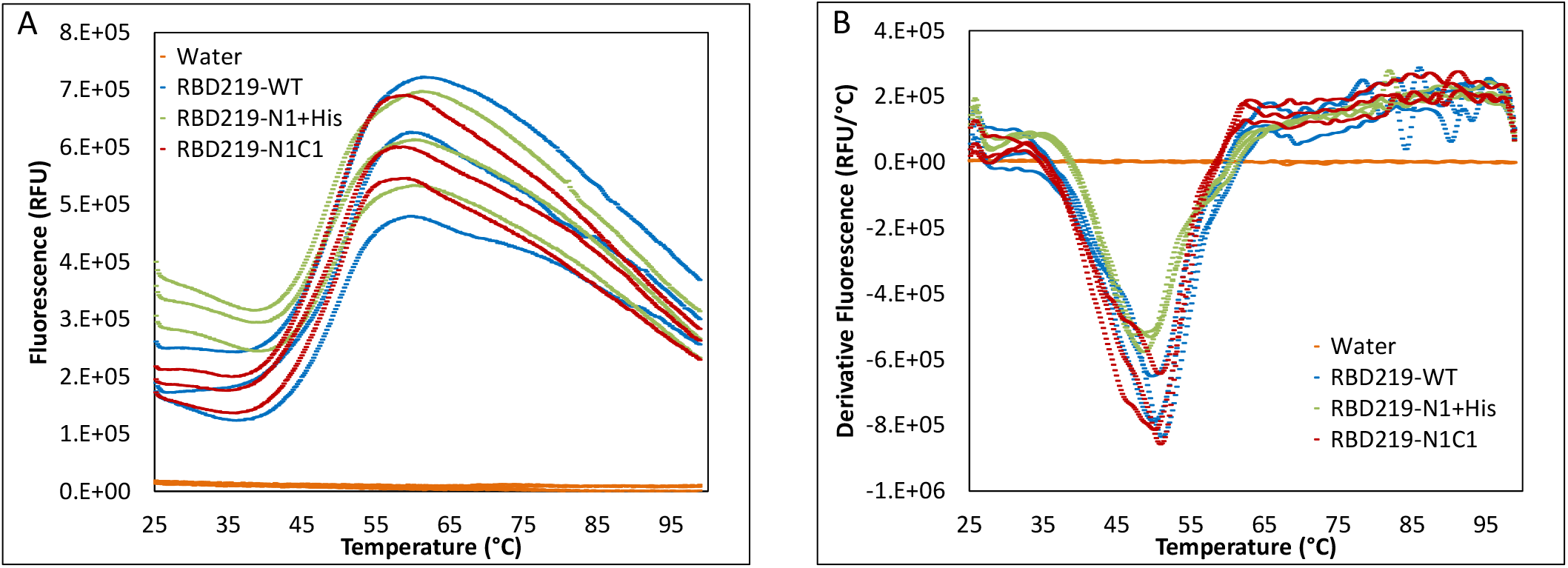
Thermal shift assay for RBD219-WT, RBD219-N1+His, and RBD219-N1C1. (A) the fluorescence-temperature plot and (B) the derivative fluorescence-temperature plot.

### 3.6. *in vitro* Functionality comparison of the RBD proteins in an ACE-2 binding study

To examine the *in vitro* functionality of the purified RBDs, we evaluated their ability to bind to a recombinant ACE-2 protein reflecting the human receptor, ACE-2 (Figure 7). No significant binding was found between ACE-2 and BSA, indicating that RBD binding to ACE-2 in this assay was specific. No statistical difference (p=0.935) among the binding of ACE-2 to RBD219-WT, RBD219-N1+His, or RBD219-N1C1 proteins was seen, indicating that hyperglycosylation on RBD219-WT, the N1 deletion, or the cysteine mutation on RBD219-N1C1 did not impact the in vitro ability of these proteins to bind to the ACE-2 receptor.

**Figure 7.**
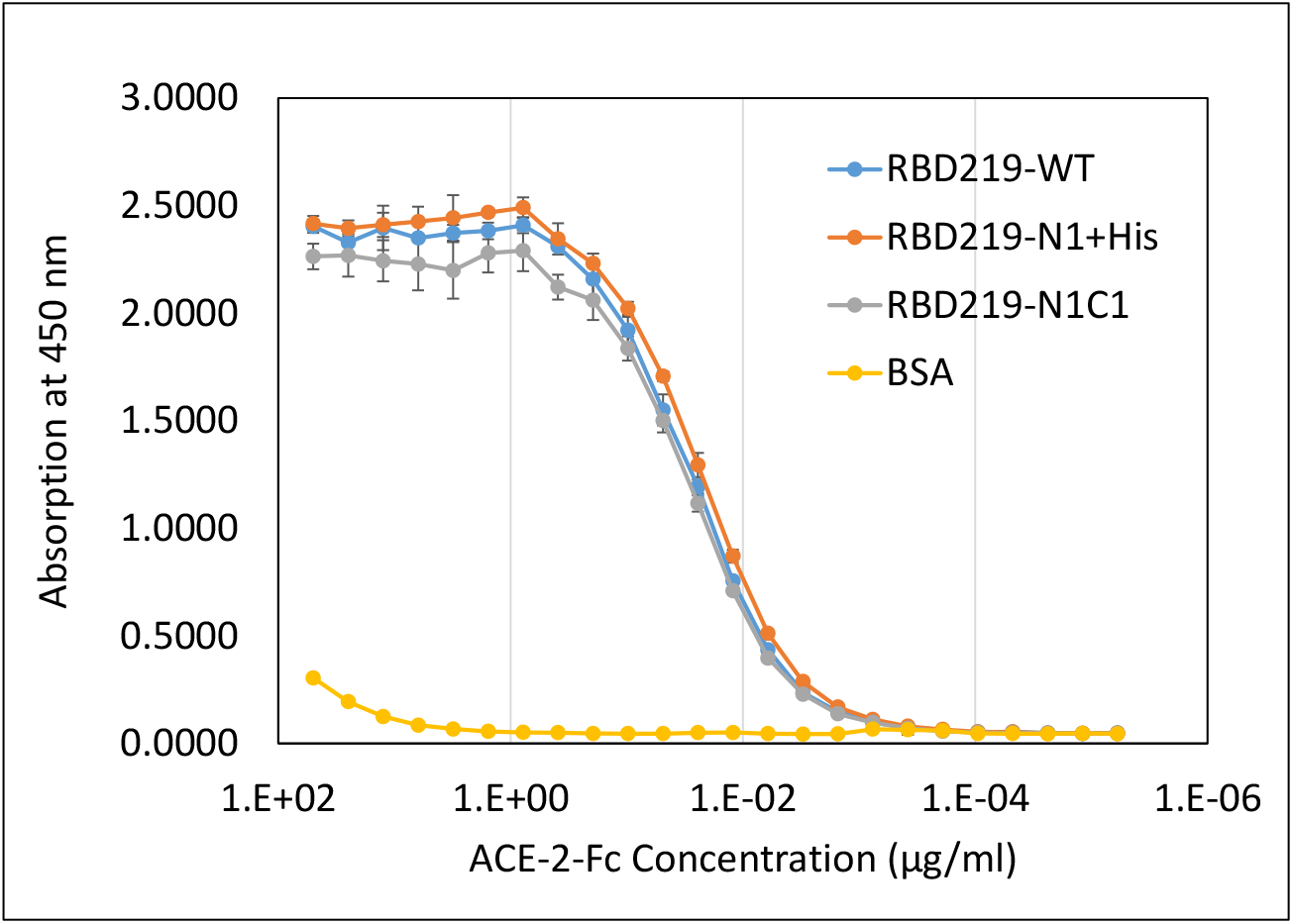
ACE-2 binding study of RBD219-WT, RBD219-N1+His, and RBD219-N1C1.

## 4. DISCUSSION

As COVID-19 spreads worldwide, it has now become predominant among populations living in poverty, especially in crowded urban areas and mega-cities located in low- and middle-income countries (LMICs) [30]; take India and Brazil as examples, the majority of the cases, over 2.1 million and 1.9 million, are reported in Maharashtra state, where Mumbai city is located, and State of Sao Paulo, where Sao Paulo city is located, respectively [1]. In the race to develop a vaccine against COVID-19, many candidates have moved forward to clinical trials. Based on several COVID-19 clinical trials databases and the World Health Organization (WHO) tracker, there are 63 vaccines currently in Phase 1 to 3 clinical trials [31, 32].

Among them, 17 are recombinant subunit vaccines, mainly using either recombinant spike or RBD proteins as a vaccine antigen [31, 33]. Within these recombinant subunit vaccine candidates, two vaccines, spike trimer from Novavax [34-36] and RBD dimer from Anhui Zhifei Longcom Biophamacetical [37, 38], are currently in Phase 3 and fifteen vaccines are in Phase 1/2, including the RBD219-N1C1 candidate presented in this manuscript [26, 32]. However, it is noted that many of these recombinant protein-based vaccine candidates are expressed using mammalian cell or insect cell systems [20, 34, 37, 39, 40] that are not always considered the most cost-effective and globally accessible.

To develop a safe and effective vaccine that is also affordable and easily accessible, we have focused our vaccine development efforts using a yeast-based expression system, similar to the low-cost recombinant hepatitis B vaccine expressed in yeast that is produced in several LMICs, including Brazil, Cuba, India, Indonesia, and elsewhere [26]. In this study, we, therefore, evaluated three different yeast-expressed antigen constructs, RBD219-WT, RBD219-N1, and RBD219-N1C1 by comparing their fermentation yield, biophysical characteristics, and *in vitro* functionality.

During the initial expression of RBD219-WT, it was found that the protein was hyperglycosylated, which could pose challenges during the production process and impact yields, reproducibility, and stability. Based on the lessons learned from the development of the yeast-expressed SARS-CoV RBD [4], we removed the N-331 residue, generating RBD219-N1. This increased the uniformity of the RBD size by SDS-PAGE (Supplementary Figure S1). However, we discovered that dimer formation via intermolecular disulfide bridging occurred during expression of the RBD219-N1 protein. While Dai et al. have recently shown that an RBD dimer improved immunogenicity of their vaccine candidate [41], we note that dimer formation is typically considered a challenge while establishing production reproducibility, scalability, stability and quality characteristics in support of regulatory enabling documentation. When investigating the structure of RBD219-N1 using PDB ID 6XEY [42], we saw nine cysteine residues, eight of which formed disulfide bonds (Supplementary Figure S2) while one, C538, was free to form intermolecular disulfide bonds. In RBD219-N1C1, we therefore mutated the free C538 residue to alanine to avoid this issue. That residue has not been shown to be an epitope involved in triggering neutralizing antibodies.

Densitometry of Coomassie blue-stained SDS-PAGE gels is generally used to evaluate the protein yield during the fermentation runs [13, 43, 44]; however, in this case, due to the high levels of likely HCP impurities observed in the FS for both RBD219-WT and RBD219-N1, the method could not be applied, and hence, we had to use western blots to assess the yields of the unpurified proteins after fermentation. By using this immunoblot-based method, we were able to quantify the protein production in the fermentation supernatants without the interference of HCPs. The protein yield quantified using western blot indicated that in the FS, RBD219-N1C1 had the highest yield among the three constructs. Due to the high HCP background found in RBD219-WT FS, QXL was used to first capture most of the HCPs leaving the target protein in the flow-through, this flow-through was further purified using Butyl HP followed by SEC. As for the RBD219-N1 construct, the target protein was barely visible in the FS when analyzed by SDS-PAGE, and thus, we were unable to purify tag-free RBD219-N1 protein; instead, a hexahistidine-tagged protein was expressed and purified by metal affinity column. RBD219-N1C1, however, was the dominant protein found in the FS and could be easily purified by a simple two-step purification scheme similar to the yeast expressed SARS-CoV RBD219-N1 [13]. Therefore, RBD219-N1C1 was selected for further process development and scale-up.

While further evaluating the purified RBDs by SDS-PAGE and western blot, dimers were observed in purified RBD219-WT and RBD219-N1+His only under the non-reduced condition, suggesting potential intermolecular disulfide bond formation. Mutating the free cysteine residue successfully reduced the propensity of oligomerization during fermentation, as no dimer formation was found in purified RBD219-N1C1. Additionally, as part of the stability evaluation, when monitoring the size using DLS, purified RBD219-N1C1 remained at 30 kDa for approximately 30 days while RBD219-WT and RBD219-N1+His continued to form oligomers. However, when determining the diffusion interaction parameter (K_D_) using DLS and assessing surface hydrophobicity (S_0_) using extrinsic fluorescence, we observed that RBD219-WT and RBD219-N1C1 shared similar K_D_ and S_0_ values while RBD219-N1+His showed a lower K_D_ and higher S_0_ values, implying a higher tendency to form oligomers. The high oligomerization tendency for RBD219-N1+His is likely due to the additional hexahistidine as we also performed an extrinsic fluorescence study on a hexahistidine-tagged version of RBD219-WT (RBD219-WT+His), and the results indicated a much higher surface hydrophobicity value (12.7 RFU*L/mg; supplementary Figure S3) than RBD219-WT. The pKa of the imidazole ring in histidine is approximately 6.0, suggesting that at pH 7.5, this aromatic ring is likely non-protonated, which makes this additional hexahistidine more hydrophobic.

When assessing the secondary and tertiary structures, far-UV circular dichroism spectra revealed that RBD219-WT, RBD219-N1+His, and RBD219-N1C1 purified proteins had similar secondary structures. The melting temperatures evaluated by CD and thermal shift assays also indicated that these three RBDs shared similar thermal stability for both secondary and tertiary structures. Additionally, Li *et al*. have derived neutralizing monoclonal antibodies from COVID-19 patients, and the most potent neutralizing antibodies that recognized the RBD blocked ACE-2 binding [42]. Thus, confirming the ACE-2 binding is crucial to ensure the epitopes within the ACE-2 binding site are still intact. A similar binding affinity to ACE-2 among all three RBDs was observed, suggesting that removing the first amino acid and/or mutating the free cysteine to alanine did not impact protein structure or functionality.

While the yeast-expressed RBD can be a potent vaccine candidate, we also recognize potential limitations and concerns: (1) N-glycosylation: Even though the ACE-2 binding site contains potent neutralizing epitopes, several recent studies indicated that some neutralizing antibodies recognize epitopes outside of the binding site, such near N343 [45, 46]. Yeast-derived glycans typically contain high levels of mannose [47], while mammalian cell-derived glycans are more complex. In the case of mammalian-cell culture expressed SARS-CoV-2, the N343 residue is highly fucosylated [48], and thus, antibodies raised against the yeast-derived protein might have a different quality. Notably, yeast-derived mannosylation has been shown to induce mannose receptor-mediated macrophage recruitment that might actually enhance immunogenicity [49, 50]. In a preclinical study conducted in mice, the yeast-expressed RBD was indeed able to induce high levels of antigen-specific antibodies and neutralizing antibodies [25].

(2) Design: The C538A mutation was designed to prevent dimer formation via intermolecular disulfide bond bridging and to improve stability. Since this mutation did not occur naturally, it could potentially affect immunogenicity, efficacy and safety of the vaccine antigen. However, the preclinical data indicated that RBD219-WT and RBD219-N1C1 were able to induce similar levels of neutralizing antibodies [25]. Since it is unknown whether the C538A mutation may have a safety effect, it is important this is monitored in the clinical settings. Nevertheless, as C539 is not conformationally close to the RBM (Supplementary Figure S2), and this cysteine was naturally forming a disulfide bridge with C590 (PDB ID: 6XEY) [42], the area near this residue is less likely to trigger neutralizing antibodies. Ongoing studies are evaluating a shortened RBD sequence that excludes C539 at the C terminus to preserve the native sequence.

(3) Antigen length: Studies have demonstrated that the neutralizing epitopes of the SARS-CoV-2 S protein are located in the N-terminal domain and the RBD of S1, and potentially on S2 [42, 51]. During the transmission among hosts, viral mutations can occur. Knowing that the RBD is only part of the spike protein, it may be more vulnerable to viral escape mutations, especially when the mutations are observed in the RBM region. Based on the lessons learned from SARS, it was discovered that some of the neutralizing antibodies recognizing the RBD of SARS-CoV strains from the first outbreak (Urbani, Tor2) did not possess potent neutralization ability against strains isolated from the second outbreak (GD03) [52]. The recently emerging SARS-CoV-2 South Africa variant (B.1.351) contains a concerning mutation, E484K, located in the RBM, which reduced the neutralizing ability of monoclonal antibodies and human convalescent sera raised against earlier SARS-CoV-2 variants [53, 54] and also negatively impacted the efficacy of several full-length S-protein vaccines [55]. Studies are underway to elucidate this question for RBD219-N1C1, and may also include the expansion of the antigen sequence beyond the RBD.

## 5. Conclusions

In this study, we have modified genetically and generated three different SARS-CoV-2 RBD constructs. The RBD219-N1C1 construct provided higher protein yields in the FS (∼280 mg RBD219-N1C1/L of FS), and the fermentation process was less impacted based on the detection of fewer impurities. This facilitated the use of a straightforward and relatively efficient initial purification scheme, with an approximate recovery yield of ∼189 mg of purified RBD219-N1C1/L of FS. With respect to biophysical characteristics, unlike RBD219-WT and RBD219-N1 proteins, the purified RBD219-N1C1 protein does not appear to form dimers via intermolecular disulfide bridging. Additionally, RBD219-WT, RBD219-N1+His, and RBD219-N1C1 showed similar thermal stability and the same binding affinity to their receptor, ACE-2, further suggesting that the deletion of the first amino acid and mutation of the free cysteine did not cause any significant structural changes. Based on all the data, we conclude that the RBD219-N1C1 construct is the superior candidate to move forward. For this vaccine candidate to advance to clinical development, we have extensively studied and developed a robust, scalable, and reproducible production process. This construct and its initial process for protein production have been transferred to an industrial manufacturer, who has successfully produced and scaled it and has now advanced this vaccine candidate into a Phase I/2 clinical trial [26].

## ABBREVIATIONS

COVID-19: Coronavirus disease 2019
SARS: severe acute respiratory syndrome
CoV: coronavirus
S: spike
RBD: receptor-binding domain
DO: dissolved oxygen
FS: fermentation supernatant
CV: column volume
%CV: coefficient of variation
DLS: dynamic light scattering
CD: circular dichroism
ACE-2: angiotensin-converting enzyme 2.

## Author contributions

WHC conceived the study, designed and performed experiments, interpreted data, and wrote the manuscript; JW performed experiments and interpreted data; RTK, RA, ZL, JL, LV, CP, BK, MJV, ACAL, and JAR performed experiments; JP participated in interpreting data, reviewed/edited the manuscript; PMG reviewed/edited the manuscript ad effected logistic and supervision; US reviewed/edited the manuscript; BZ reviewed/edited the manuscript; PJH reviewed/edited the manuscript; MEB reviewed/edited the manuscript US, BZ, PJH, and MEB also provided scientific guidance on the project.

## Acknowledgments

This work was supported by the Robert J. Kleberg Jr. and Helen C. Kleberg Foundation; Fifth Generation, Inc. (Tito’s Handmade Vodka); JPB Foundation, NIH-NIAID (AI14087201); and Texas Children’s Hospital Center for Vaccine Development Intramural Funds. We also would like to thank PATH Center for Vaccine Innovation and Access (Seattle, WA, USA) for their guidance as well as technical and intellectual support.

