## Supplementary for "Genetic Modification to Design a Stable Yeast-expressed Recombinant SARS-CoV-2 Receptor Binding Domain as a COVID-19 Vaccine Candidate"

| **RBD219-WT** | **RBD219-N1 (+/- His)** | **RBD219-N1C1** |
| --- | --- | --- |
| A  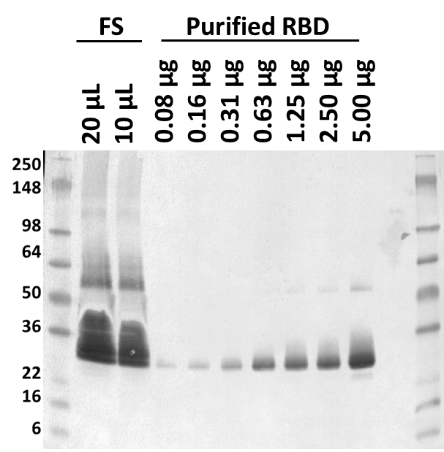 | E  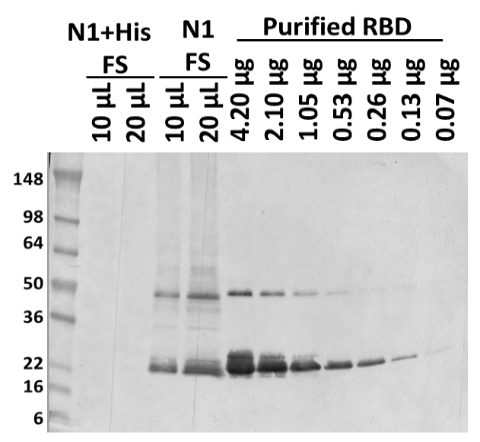 | I  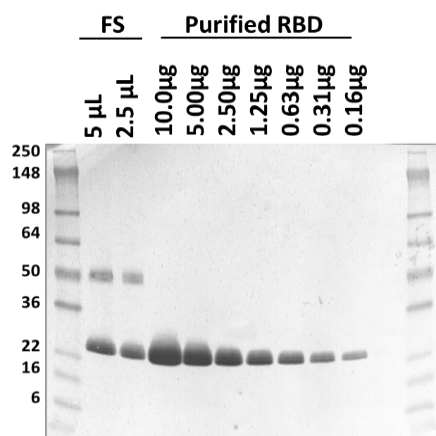 |
| B | F | J |
| C  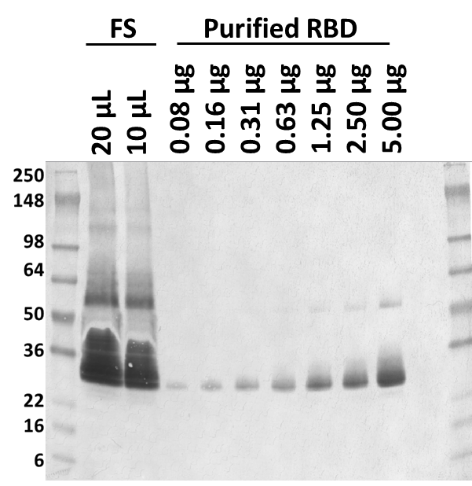 | G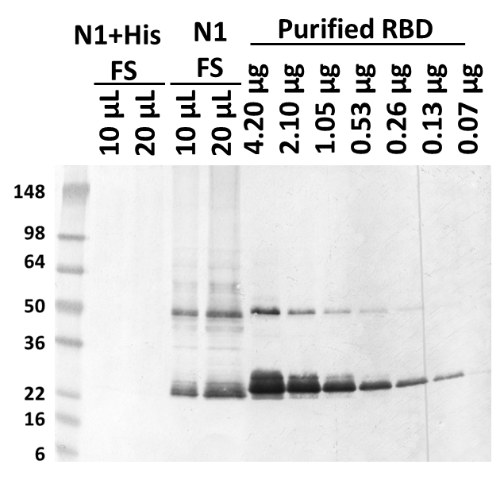 | K  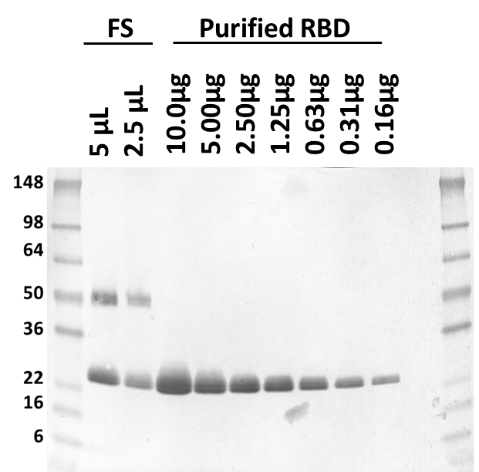 |
| D | H | L |

| Log (intensity) | Log (loaded weight of RBD) |
| --- | --- |
| 3.35211 | 0.69897 |
| 3.12652 | 0.39794 |
| 3.06448 | 0.09691 |
| 2.9821 | -0.20412 |
| 2.77403 | -0.50515 |
| 2.56677 | -0.80618 |
| 2.46465 | -1.10721 |

Log (loaded weight of RBD)

| Log (intensity) | N1+His |
| --- | --- |
| 2.5163 | -1.18293 |
| 2.83504 | -0.881901 |
| 3.06115 | -0.580871 |
| 3.18624 | -0.279841 |
| 3.40572 | 0.0211893 |
| 3.58166 | 0.322219 |
| 3.67832 | 0.623249 |

Log (loaded weight of RBd)

| Log (intensity) | N1C1 |
| --- | --- |
| 2.78596 | -0.80618 |
| 2.94008 | -0.50515 |
| 3.08145 | -0.20412 |
| 3.18378 | 0.09691 |
| 3.2487 | 0.39794 |
| 3.31037 | 0.69897 |
| 3.44148 | 1 |

| Log (intensity) | Log (loaded weight of RBD) |
| --- | --- |
| 3.3421 | 0.69897 |
| 3.11193 | 0.39794 |
| 3.0534 | 0.09691 |
| 2.88259 | -0.20412 |
| 2.75194 | -0.50515 |
| 2.61166 | -0.80618 |
| 2.48552 | -1.10721 |

Log (loaded weight of RBD)

| Log (intensity) | N1+His |
| --- | --- |
| 2.47022 | -1.18293 |
| 2.66552 | -0.881901 |
| 2.98832 | -0.580871 |
| 3.19534 | -0.279841 |
| 3.38465 | 0.0211893 |
| 3.55853 | 0.322219 |
| 3.73988 | 0.623249 |

Log (loaded weight of RBD)

| Log (intensity) | N1C1 |
| --- | --- |
| 2.78596 | -0.80618 |
| 2.95316 | -0.50515 |
| 3.0983 | -0.20412 |
| 3.19549 | 0.09691 |
| 3.26812 | 0.39794 |
| 3.31983 | 0.69897 |
| 3.44255 | 1 |

**Figure S1** Western blot of RBD219-WT fermentation supernatant and purified RBD219-WT (A, C) and the standard curve generated using the purified RBD219-WT (B, D). Western blot of RBD219-N1 and N1+His fermentation supernatant and purified RBD219-N1+His (E, G) and the standard curve generated using the purified RBD219-N1+His (F, H). Western blot of RBD219-N1C1 fermentation supernatant and purified RBD219-N1C1 (I, K) and the standard curve generated using the purified RBD219-N1C1 (J, L)

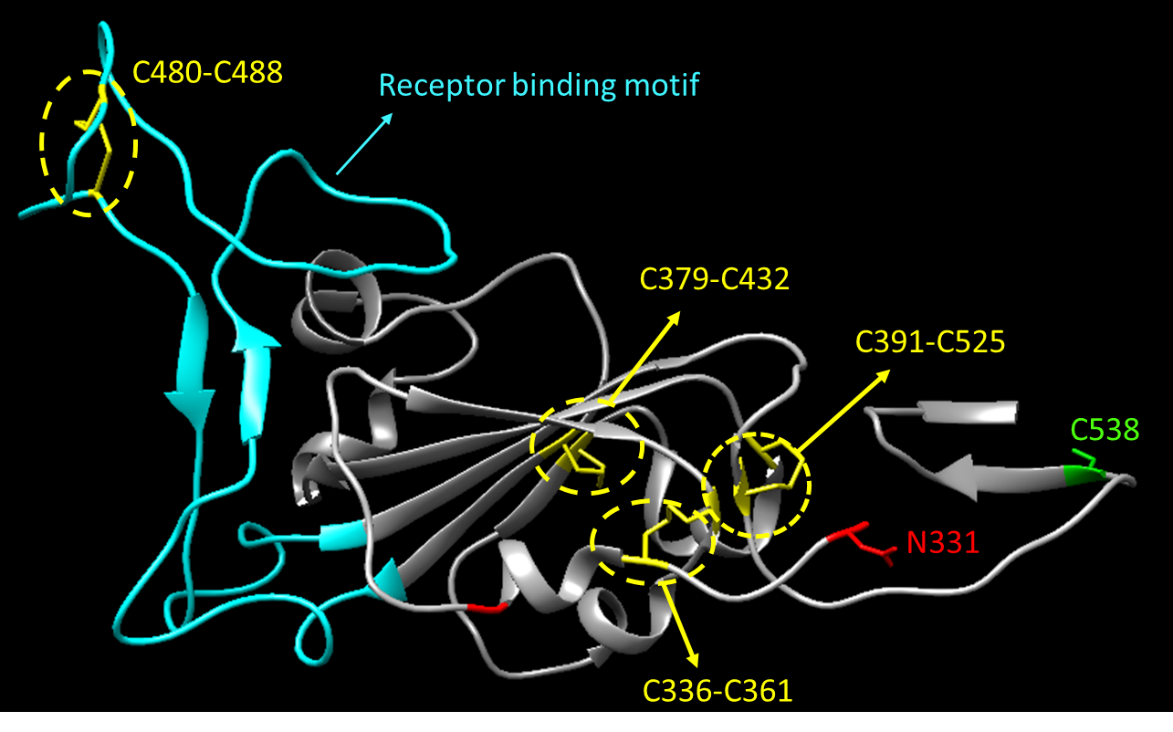

**Figure S2.** The structure of RBD219-WT is extracted from 6XEY using Chimera 1.14. The four disulfide bond formation at C336/C361, C379/C432, C391/525, and C480/488 are highlighted in yellow and the free cysteine is highlighted in green. The two glycosylation sites N331 and N343 are highlighted in red. The receptor-binding motif (RBM) is highlighted in cyan.

Fluorescence intensity (RFU)

| Concentration (mg/mL) | RBD21-WT+His | RBD219-WT | RBD219-N1+His | RBD219-N1C1 |
| --- | --- | --- | --- | --- |
| 1 | 15817 | 4974 | 12892 | 4829 |
| 0.5 | 9284.5 | 3564 | 7254 | 4197.5 |
| 0.25 | 6260 | 2583 | 5131.5 | 2903 |
| 0.125 | 4853 | 2104.5 | 4053.5 | 2531 |
| 0.0625 | 4491 | 2045 | 3667.5 | 2478.5 |
| 0.03125 | 3279 | 1967.5 | 3080.5 | 2304 |
| 0.015625 | 3264 | 1969.5 | 2966.5 | 2190 |
| 0.0078125 | 2841.5 | 1944.5 | 3063.5 | 2349 |

**Figure S3** Fluorescence intensity vs. concentration plot for different RBD variants
